# ACKR5/GPR182 is a scavenger receptor for the atypical chemokine CXCL17, GPR15L and various endogenous peptides

**DOI:** 10.1101/2024.06.01.596940

**Authors:** Max Meyrath, Christie B. Palmer, Jean-Marc Plesseria, Joy Darcis, Desislava Nesheva, Marcus Thelen, Daniel F. Legler, Rob Leurs, Julien Hanson, Martyna Szpakowska, Andy Chevigné

## Abstract

GPR182/ACKR5, the most recently deorphanized chemokine receptor, is mainly expressed on endothelial cells and was proposed to act as a scavenger regulating the availability of a large set of chemokines. In this study, we first established the exact profiles and ranking of the chemokines binding to the human and mouse GPR182. We confirmed the high promiscuity of GPR182 towards XC, CC and CXC chemokines and a clear difference in the chemokine repertoires of the human and mouse orthologues. We next demonstrated that, beyond classical chemokines, GPR182 exhibits potent binding to the chemoattractant protein GPR15L/C10orf99, the atypical chemokine CXCL17 and various endogenous peptides, mainly from the opioid, apelin, and PACAP families. We also showed that these newly identified ligands engage GPR182 through varied binding modes. While GPR15L, just like classical chemokines, predominantly engages GPR182 via its N terminus, conversely to the C terminus-dependent binding to its cognate receptor GPR15, CXCL17 exhibits a more complex interaction, relying on both the N and C terminus. The binding mode of the newly identified peptide ligands also differ from the interactions with their cognate receptors. Our findings establish the first scavenger receptor for CXCL17 and GPR15L and advance the understanding of GPR182 ligand interactions, suggesting a regulatory role beyond chemokines.

## Short report

Over the past years, atypical chemokine receptors (ACKRs) have emerged as important regulators of chemokine functions. ACKRs, unable to trigger G protein-dependent signalling, sequester or internalize chemokines, thereby shaping their gradient and regulating their availability to the respective classical chemokine receptors. Four ACKRs have been identified so far, but recent reports suggest that GPR182, an orphan receptor expressed on endothelial cells, which shares about 30% of sequence similarity with its closest relative ACKR3, could serve as an atypical chemokine receptor (ACKR5) for a broad spectrum of chemokines from different families^1-3^. Although discrepancies on its exact ligand selectivity persist, studies have consistently demonstrated a strong constitutive interaction with β-arrestin with, unlike other ACKRs, no ligand-induced effect, rendering its investigation challenging^1,3^. It was further suggested that GPR182 may have an unconventional chemokine interaction mode, relying on the glycosaminoglycan (GAG)-binding motif, but this hypothesis requires further validation^2^. Furthermore, whether GPR182, similarly to ACKR3, can also bind non-chemokine ligands remains an open question^4^.

In this study, we first established a comprehensive chemokine repertoire for GPR182 by performing systematic binding competition studies with fluorescently labelled CXCL12, allowing to precisely determine and compare pIC_50_ values (**Figures 1a and S1**). We identified CXCL13 (pIC_50_ = 7.67), CXCL12 (pIC_50_ = 7.24), CCL19 (pIC_50_ = 7.21) and CCL25 (pIC_50_ = 7.17) as the strongest binders, followed by other chemokines from the CC, CXC and XC families: CXCL10, the orphan chemokine CXCL14, CXCL16, XCL1, CXCL2, CXCL3, CXCL11, CXCL9, CCL24, CCL1, CCL21, CCL28, CCL20, CCL26 and CCL11 (**Figure 1a**). Similar potencies were obtained in binding competition studies with a labelled CC chemokine, CCL19 (**Figure S2a**). In addition, we showed that the viral macrophage inflammatory protein II (vMIP-II or vCCL2), a broad-spectrum CC chemokine encoded by the human herpesvirus 8 (HHV-8), is a potent binder of the human GPR182 (pIC_50_ = 7.87) and a weak binder of the mouse GPR182 (pIC_50_ = 6.13). We indeed confirmed that several chemokines exhibited clear differences in potencies towards the human and mouse GPR182 orthologues, notably CCL19, CCL22 and CCL27 (**Figure 1a and S3**)^4,5^. These data validate GPR182 as a broad-spectrum receptor of CC, CXC and XC chemokines and establish a precise ranking of both the mouse and human chemokines on their respective receptors^1-3,5^.

**Figure 1.**
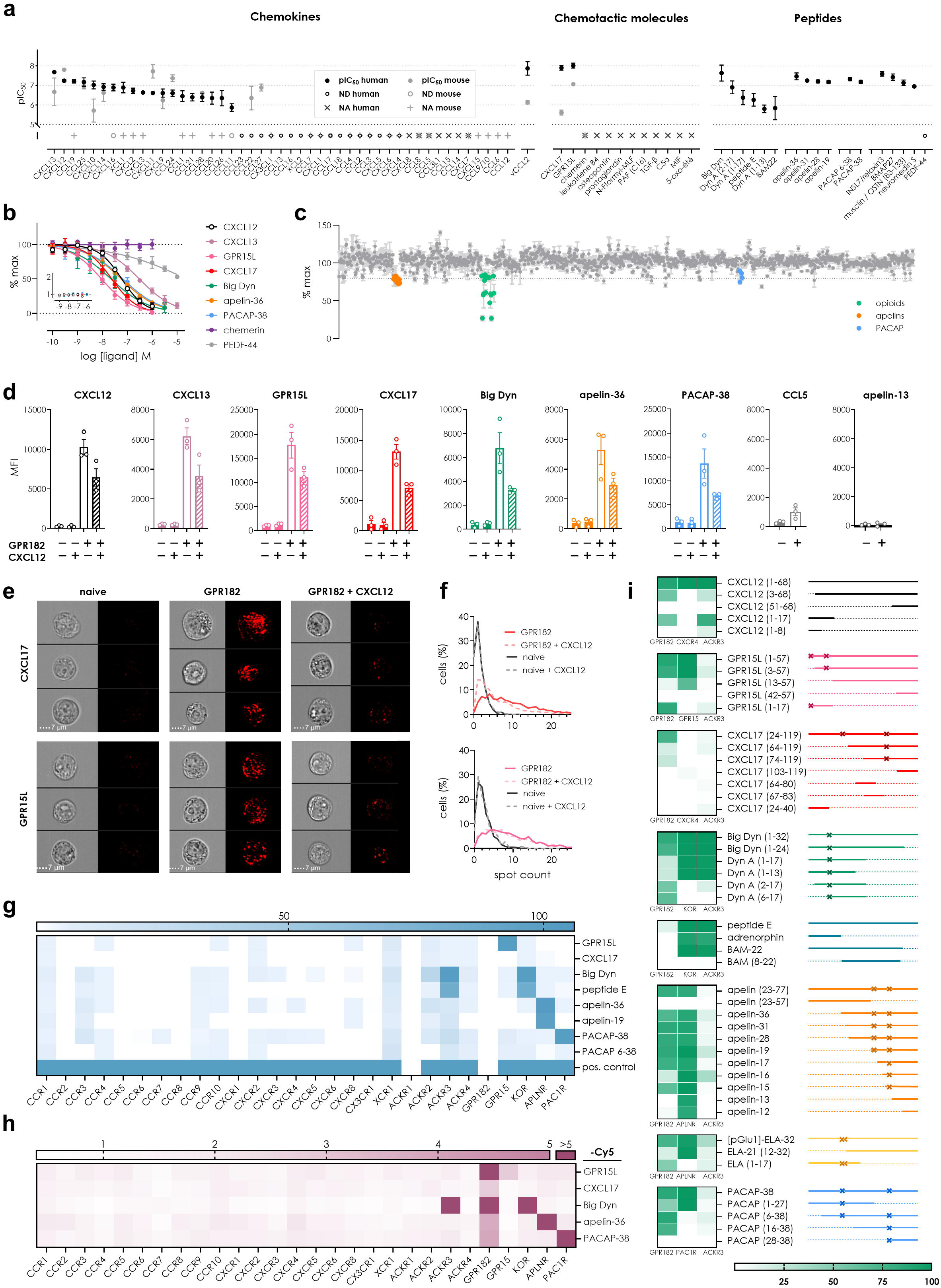
GPR182 selectively binds and internalizes several chemokine and non-chemokine ligands from different families. **(a)** pIC_50_ values of all human and mouse chemokines (left), a set of other chemotactic molecules (middle) and hits from a peptide library screen determined by flow cytometry binding competition with human or mouse AZ647-labelled CXCL12 (10 nM) on human (black) and mouse (grey) GPR182 orthologues, respectively. ND: pIC_50_ value not determinable; NA: no competition with CXCL12 in the concentration range tested (up to 1 µM). **(b)** Full concentration– response curves of selected ligands from different families determined by flow cytometry binding competition with AZ647-labelled CXCL12 (10 nM) on GPR182. Inset: Ligand-induced β-arrestin 1 recruitment to GPR182 expressed as fold to vehicle determined by NanoBiT. **(c)** Screen of three peptide libraries containing over 1300 peptides tested at 1 µM in NanoBRET binding competition with AZ568-labelled CXCL12 (30 nM). Families of interest are highlighted in colour. **(d)** GPR182-mediated uptake of Cy5-labelled chemokines (10 nM) or peptides (20 nM) in competition with unlabelled CXCL12 (300 nM) visualized in naïve or HEK293-GPR182 cells by imaging flow cytometry and expressed as mean fluorescence intensity (MFI). **(e)** Uptake of Cy5-labelled CXCL17 (top) or GPR15L (bottom) and **(f)** corresponding percentage of cells with a given number of distinguishable vesicle-like structures (spots). Scale bar: 7 µM. **(g)** Agonist activity of representative new GPR182 ligands towards all chemokine receptors and cognate receptors evaluated in a β-arrestin 1 recruitment assay. CXCL17 and GPR15L (100 nM), other ligands (1 µM). Results are expressed as percentage of signal monitored with a positive control chemokine (200 nM) or peptide (1 µM) (**h**). Uptake of Cy5-labelled ligand uptake monitored by flow cytometry. CXCL17 and GPR15L (10 nM), other ligands (20 nM). (**g**) Results are expressed as fold change over non-transfected HEK cells. (**i**) Comparison of the efficacy of truncated ligands (1 μM) in NanoBRET binding competition (GPR182) or NanoBiT β-arrestin 1 recruitment (cognate receptors or ACKR3), expressed as percentage of maximal signal (*left*). The fragment sizes are indicated and clusters of basic residues (at least 3 residues out of four) are highlighted with an × (*right*). Results are expressed as mean ± SEM of three (a, b and d) or two (c) independent experiments. For g, h and i, results are expressed as mean of three independent experiments. e and f are representative data from one out of three experiments depicted in d.

Considering the extended spectrum of chemokines recognized by GPR182, we wondered whether it could be a receptor for a broader variety of ligands. We therefore first screened an array of non-chemokine chemotactic mediators, including molecules such as leukotriene B4 (LTB4), formyl peptides but also small soluble proteins such as chemerin, the chemoattractant proteins GPR15L, macrophage migration inhibitory factor (MIF), the complement component 5a (C5a) and CXCL17, an atypical chemokine strongly binding to GAGs that had not been considered in the panels tested in previous studies^6^.

While most of the chemotactic molecules showed no binding to GPR182, GPR15L (pIC_50_ = 8.00), previously demonstrated to be a low-affinity agonist for the G protein-coupled receptor GPR15^7,8^, and CXCL17 (pIC_50_ = 7.89) interacted with GPR182 with stronger potencies than any classical chemokines tested (**Fig 1a–b, Fig S2a–d**). mGPR15L, which shares 52% sequence identity with its human counterpart, bound to mGPR182 with a tenfold weaker potency (pIC_50_ = 7.06), while mCXCL17 showed poor binding (pIC_50_ = 5.61) (**Figure S2d**). GPR15L recently garnered attention for its direct chemotactic role in the immune function in the skin and mucosa. Interestingly, although it shares some features with CC chemokines, including a conserved N-terminal CC motif, GPR15L does not match the criteria to be considered a chemokine, as the pairings of its two stabilizing disulfide bridges clearly differ from those found in classical chemokines^7^. Similarly, although officially classified as a chemokine, CXCL17 displays fundamental structural differences compared to conventional chemokines, does not bear the usually positioned N-terminal cysteine motif and is predicted not to adopt the typical chemokine fold^6^. Attempts to identify the receptor for CXCL17 remain so far unsuccessful. Recent reports, however, suggested that CXCL17 is a low-affinity agonist for the Mas-related GPR family member X2 (MRGPRX2), a low-selectivity receptor for peptides, and that it could act as an allosteric inhibitor of CXCR4^9,10^. This study establishes GPR182 as a high-affinity receptor also for two biologically active small non-chemokine proteins, GPR15L and CXCL17.

These results prompted us to further explore the binding repertoire of GPR182 and examine whether, similar to ACKR3, GPR182 could also bind short endogenous peptides. Using a NanoBRET assay developed for high-throughput binding competition studies with fluorescently labelled CXCL12, we screened libraries combining over 1300 bioactive peptides and analogues, covering a wide variety of ligand families, including 150 orphan peptides (**Figure 1c**). We were able to identify several GPR182 binders clustering into three distinct families: the opioid peptides, especially dynorphins (big dynorphin 1-32, dynorphin 2-17 and 6-17) involved in the regulation of emotional behaviours, the apelins (apelin-36, - 31, -28, -19, -17) that among other functions control blood pressure and angiogenesis and the pituitary adenylate cyclase-activating polypeptides, also known as PACAPs (PACAP-38 and 6-38), that have pleiotropic regulatory activity including in stress response. We confirmed these hits using flow cytometry binding competition on GPR182 and found that the pIC_50_ values of several of these peptides were in the same order of magnitude as most of the chemokines binding to GPR182 (**Figures 1a-b and S2e, Table S1**).

We also showed that, just like for chemokines, the binding of these new ligands to GPR182 did not induce an increase of the interaction with β-arrestins or trigger G protein activation (**Figures 1b inset and S4**). Next, to establish whether GPR182 can serve a comparable scavenger role for these new ligands as it does for chemokines, we used imaging flow cytometry to measure the uptake of labelled CXCL17, GPR15L and representatives of the different peptide families, *i*.*e*. big dynorphin, apelin-36 and PACAP-38, by GPR182-expressing cells (**Figure S5)**. We observed that, similarly to CXCL12 and CXCL13, all the newly identified ligands accumulated in vesicle-like structures exclusively in cells expressing GPR182, which was impaired in the presence of competing unlabelled CXCL12 (**Figures 1d-f and S6)**. We also showed that besides ACKR3, known to interact with opioid peptides^4^, none of the other 22 chemokine receptors was responsive to the novel ligands of GPR182, as demonstrated in β-arrestin recruitment (**Figure 1g**) and flow cytometry ligand uptake (**Figure 1h**) assays.

Chemokines typically activate their receptors through the binding of their flexible N terminus penetrating the receptor transmembrane cavity. In contrast, the receptor-activating determinants of other chemotactic molecules, such as chemerin, C5a or GPR15L, are mainly located in their C terminus. Considering the great differences in terms of size, structure and sequences of the newly identified GPR182 ligands and the previously proposed mechanism whereby the receptor recognizes the GAG-binding motif of chemokines^2^, we set to investigate its ligand interaction modes in comparison to those of their known receptors. We therefore undertook a comparative structure–activity relationship (SAR) analysis using truncated ligands lacking N- and C-terminal extremities, as well as peptides covering regions relevant to their activity towards the classical receptors **(Figure 1i, S7 and S8)**. For CXCL12, we found that the interaction mode with GPR182 was similar to ACKR3, mainly relying on the chemokine N terminus and N loop. On the other hand, the newly identified ligands engage GPR182 through varied binding modes that often differ fundamentally from the way they act on their classical receptors. Indeed, while GPR15L, much like the classical chemokines, predominantly engaged GPR182 via its N terminus, CXCL17 exhibited a more complex interaction, relying on both the N and C terminus. Further supporting this contrast, N and C terminal truncations of opioid peptides had opposite effects on GPR182 compared to the kappa opioid receptor (KOR) and ACKR3.

A different interaction mode was also observed for peptides from the apelin and PACAP families. For apelin, although the overall importance of the C-terminal part of the peptide seemed to be conserved for both GPR182 and the apelin receptor (APLNR), the length of the peptides had a different impact on the strength of the interaction with the two receptors. Interestingly, in line with the previously suggested binding mode for chemokines^2^, we found that the interaction between GPR182 and most of the new ligands appeared to be severely impaired upon removal of clusters of basic amino acids. This was for example illustrated by the absence of the highly positively charged N terminus of GPR15L, the central segment of PACAP, big dynorphin or CXCL17, or the positively charged C terminus of apelin. However, a more complex or alternative binding mode may be involved for other ligands as exemplified by CXCL12, which does not show such a positively charged cluster within its N terminus (**Figures 1i, S7 and S8**).

Overall, we reveal that GPR182, similar to its closest relative ACKR3, has regulatory functions that extend far beyond the classical chemokine ligands. For most of the ligands identified in this study, their potencies towards GPR182 are similar to those of chemokines. Although the exact physiological relevance of these new pairings remains to be established, GPR182 appears to accommodate a unique ligand repertoire and unconventional promiscuous binding mode, which clearly distinguish it from other ACKRs. In light of the present study, new highly charged non-chemokine ligands are likely to be identified for GPR182 in the near future. Finally, this study also highlights the limitations and porosity of the classification of chemokines and their receptors and the existence of possible crosstalk and interplays with other ligand or receptor families (**Figure S9**).

## Material and methods

### Peptides and proteins

If not otherwise stated, non-labelled murine and human chemokines were purchased from Peprotech. hCXCL8, hCXCL9, hCXCL11, hCCL2, hCCL5, AZDye-labelled chemokines hCXCL12-AZ647, mCXCL12-AZ647, hCCL19-AZ647, hCXCL12-AZ568 were purchased from Protein Foundry, hCCL1, hCXCL17, mCXCL17 hXCL2, mCXCL7 from R&D Systems, mCXCL3 mXCL1, mCX3CL1, mCCL1 and mCCL25 from BioLegend and mCXCL15 from Bio-Connect. For comparative analysis and hit confirmation, hCXCL17 was also purchased from BioLegend and from Prospec. GPR15L, C5a and TNF-α were purchased from Peprotech, chemerin, MIF and osteopontin from R&D Systems, 5-oxo-eicosatetraenoic acid and prostaglandin from Santa Cruz, Leukotriene and PAF from Tocris. N-Formylmethionyl-leucyl-phenylalanine (fMLF), as well as hGPR15L for hit confirmation were purchased from Phoenix Pharmaceuticals. N-or C-terminally truncated peptides were custom synthesized from JPT or purchased from Phoenix Pharmaceuticals.

All fluorescent peptides and chemokines were generated using Cy-5 N-hydroxysuccinimide (NHS) ester QuickStain protein labelling kit (Amersham) according to manufacturer’s protocol.

The Chop-Suey/Variety Peptide Library (L-001, 238 peptides), Bioactive Secretory Peptide Library (L-009, 1013 peptides) and Orphan Receptor Peptide Ligand Library (L-005, 150 peptides) and selected individual peptides from the opioid, PACAP and apelin families were purchased from Phoenix Pharmaceuticals.

### Cell culture

HEK293T cells were purchased from Abcam and grown in Dulbecco’s modified Eagle medium (DMEM) supplemented with 10% fetal bovine serum (Sigma) and penicillin/streptomycin (100 Units per ml and 100 μg per ml). HEK293T cells stably expressing untagged human or mouse GPR182 (hGPR182 or mGPR182) or hCXCR4, hACKR3 or hGPR182 N-terminally fused to Nanoluciferase were established using pIRES-puromycin vectors and were grown in DMEM medium supplemented with 10% fetal bovine serum (Sigma) and penicillin/streptomycin (100 Units per ml and 100 μg per ml) and puromycin (5 μg per ml).

### Ligand binding competition monitored by flow cytometry

Ligand binding to a receptor was monitored using 2.0 × 10^5^ HEK293T stably expressing murine or human receptor GPR182. Ligands were serially diluted in the presence of 10 nM murine or

human CXCL12-AZ647 or 20 nM human CCL19-AZ647 and incubated for 2 hours with the cells at 4°C. Cells were washed with FACS buffer (PBS, 1% BSA, 0.1% NaN_3_) and incubated with Zombie Green viability dye (BioLegend) to exclude dead cells. The signal obtained from parental HEK cells was used to evaluate non-specific binding of CXCL12-AZ647 and defined as 0% binding. AZ647 signal on GPR182 expressing cells in the absence of any competitor was defined as 100% binding. Binding was quantified as mean fluorescence intensity on a NovoCyte Quanteon flow cytometer (ACEA Biosciences) using NovoExpess 1.4.1.

### Ligand binding competition monitored by NanoBRET

Ligand binding to GPR182, ACKR3 and CXCR4 was monitored by NanoBRET on living cells and isolated membranes. HEK293T cells, stably expressing the receptors, N-terminally fused to Nanoluciferase were distributed into white 384-well plates (1.5 × 10^4^ cells per well). A single concentration (1 µM, for library screening) or increasing concentrations of ligands were then added to the cells and incubated for 5 minutes on ice. Cells were subsequently incubated with CXCL12-AZ568 (2 nM for ACKR3, 30 nM for CXCR4 and GPR182) and incubated for 2 hours on ice. NanoGlo Live Cell Assay System substrate was then added and donor emission (450/8 nm BP filter) and acceptor emission (600 nm LP filter) were immediately measured on a GloMax Discover plate reader (Promega). BRET binding signal was defined as acceptor/donor ratio, and cells not treated with CXCL12-AZ568 were used to define 0% BRET binding, whereas cells that were treated with CXCL12-AZ568 alone were used to define 100% BRET binding.

### β-arrestin recruitment monitored by NanoBiT

Ligand-induced β-arrestin recruitment to receptors was monitored by Nanoluciferase complementation assay (NanoBiT, Promega). Briefly, 6 × 10^6^ HEK cells were plated in 10-cm culture dishes and 24 h later cotransfected with pNBe vectors encoding GPCRs C-terminally tagged with SmBiT and human β-arrestin-1 (arrestin-2) N-terminally fused to LgBiT. 24 h after transfection, cells were harvested, incubated 25 min at 37 °C with Nano-Glo Live Cell substrate diluted 200-fold and distributed into white 96-well plates (1 × 10^5^ cells per well). Ligand-induced β-arrestin recruitment to receptors was measured with a GloMax Discover plate reader (Promega, USA) for 20 min. Antagonist activity was measured following addition of ∼EC_80_ agonist.

For single concentration screening experiments on all chemokine receptors, the results are expressed as percentage of signal monitored with a positive control chemokine (200 nM) listed in the IUPHAR repository of chemokine receptor ligands or peptide (1 μM). For concentration– response curves, the signal recorded with a saturating concentration of full agonist for each receptor was set as 100%.

### G protein activation monitored by NanoBRET

Ligand-mediated G protein dissociation was monitored using a BRET-based G protein activity sensor based on a tricistronic plasmid, which encodes Nanoluciferase-tagged Gα_i_ subunits together with the related G_β_ and circularly permutated Venus-tagged G_γ_^11^. G protein activity was monitored through the reduction of BRET signal upon G protein subunit dissociation, as described previously^12^. In brief, 6 × 10^6^ HEK293T cells were plated in 10-cm dishes and 24 hours later co-transfected with vectors encoding G protein activity sensors and untagged receptors. 24 hours after transfection, cells were harvested and distributed into white 96-well plates (1 × 10^5^ cells/well in Opti-MEM). Next, coelenterazine-H in Opti-MEM was added to the cells followed by immediate ligand addition at indicated concentrations. BRET signal was measured with a GloMax Discover plate reader (Promega) equipped with 450/8 nm filter for donor luminescence emission and 530 nm LP filter for acceptor fluorescence emission. Ligand-induced changes in BRET ratio were expressed as fold to unstimulated condition.

### Visualization and quantification of fluorescently labelled ligand uptake by imaging flow cytometry

HEK293T cells stably expressing GPR182 or naïve HEK293T cells were distributed into 96-well plates (6.5 × 10^5^ cells/well, in Opti-MEM + 1% BSA). After 10-minute incubation with CXCL12 (300 nM) or Opti-MEM + 1% BSA, Cy5-labelled chemokines, GPR15L (10 nM) or peptides (20 nM) were added and incubated for 60 minutes at 37°C. Cells were then washed twice with FACS buffer, stained with a Zombie Green viability dye and fixed with 4.2% formaldehyde. Images of 1 × 10^4^ in-focus living single cells were acquired with an ImageStream MKII imaging flow cytometer (Amnis) using 60x magnification. Samples were analyzed using Ideas6.2 software and the number of spots per cell was determined using a mask-based software wizard.

### Quantification of fluorescently labelled ligand uptake by flow cytometry

6 × 10^6^ HEK cells were plated in 10-cm culture dishes and 24 h later cotransfected with receptors C-terminally tagged to mNeonGreen. 48 h later, they were distributed into 96-well plates (2.0 × 10^5^ cells/well, in Opti-MEM + 1% BSA) and incubated for 60 minutes at 37°C with increasing concentrations of Cy5-labelled ligands, or with a fixed concentration of chemokines (10 nM), GPR15L (10 nM) or peptides (20 nM) for selectivity screening. Cells were then washed twice with FACS buffer, stained with a Zombie UV viability dye and fixed with 4.2% formaldehyde. Uptake was quantified as mean fluorescence intensity of mNeonGreen-expressing cells subtracted by the signal obtained from non-transfected cells on a BD FACS Fortessa cytometer (BD Biosciences) using FACS Diva 8.01 (BD Biosciences)

### Curve fitting and statistical analysis

Concentration–response curves were fitted using the Hill equation with four parameters and an iterative, least-squares method (GraphPad Prism version 10.1.2). All curves were fitted to data points representing the mean of at least three independent experiments.

## Acknowledgements

This study was supported by the Luxembourg Institute of Health (LIH) through the NanoLux Platform, Luxembourg National Research Fund (INTER/FNRS CXCL12 20/15084569, PRIDE 21/16763386/CANBIO2, and AFR Opiokine 14616593), F.R.S.-FNRS-Télévie (7.8504.20, 7.4502.21 and 7.8508.22). MS and DFL received joint funding from the FNR/SNSF weave grogram (FNR CORE IMPACTT C23/BM/18068832; SNSF grant # 220358). CP, MS AC are part of the Marie Skłodowska-Curie Innovative Training Network ONCORNET2.0 “ONCOgenic Receptor Network of Excellence and Training” (MSCA-ITN-2020-ETN). The authors would like to thank Manuel Counson and Nadia Beaupain for technical and experimental support.

## Supplementary Figures

**Fig. S1.**
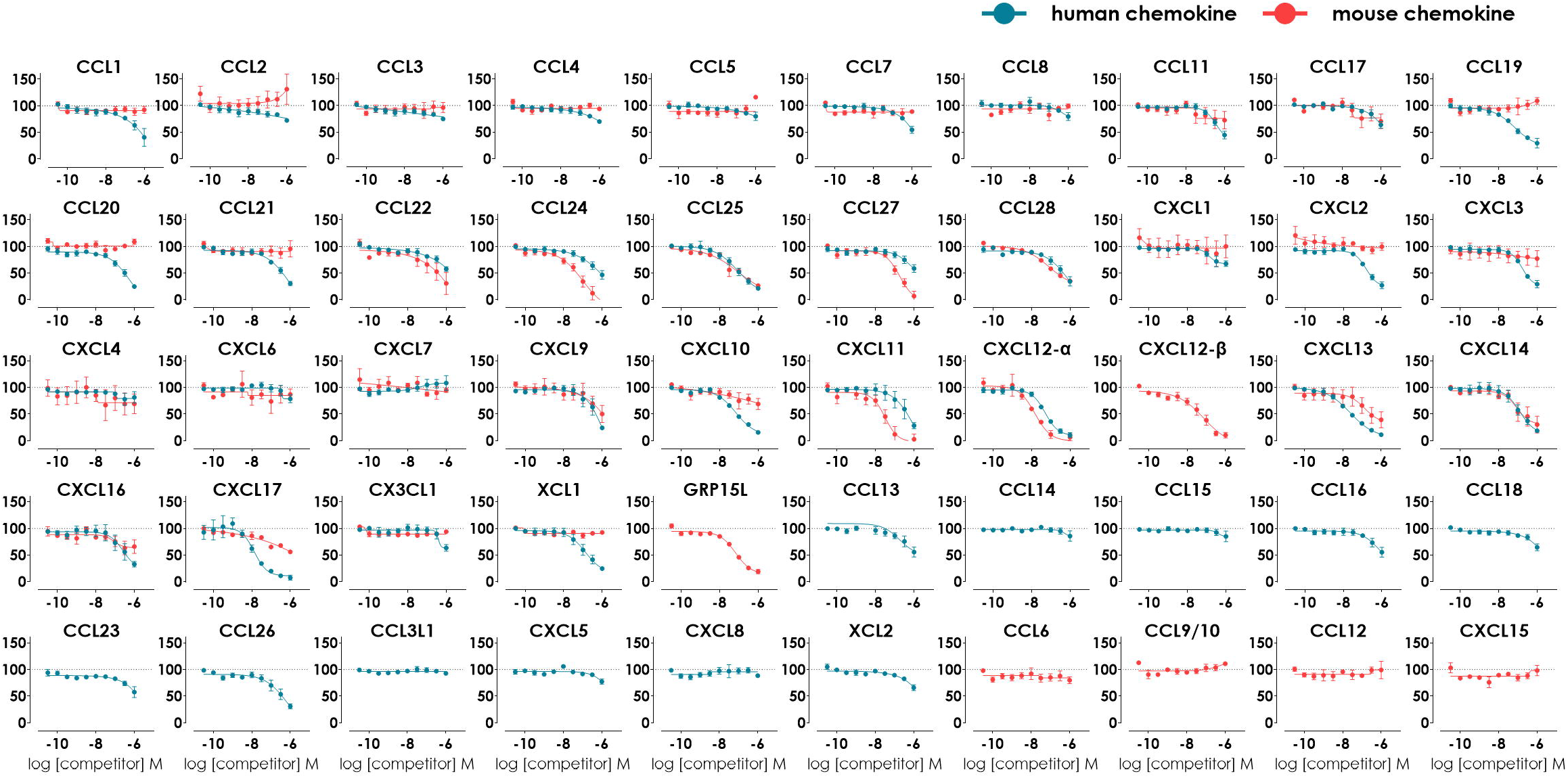
Activity screen of all the murine and human chemokines on mouse and human GPR182. Binding competition determined by flow cytometry with human or mouse AZ647-labelled CXCL12-(10 nM). Competition of human chemokines tested on hGPR182 are shown in blue, while murine chemokines tested on mGPR182 are depicted in red. Results are expressed as mean ± SEM of at least three independent experiments. pIC50 values of the individual curves were extracted and are plotted in Figure 1a.

**Fig. S2.**
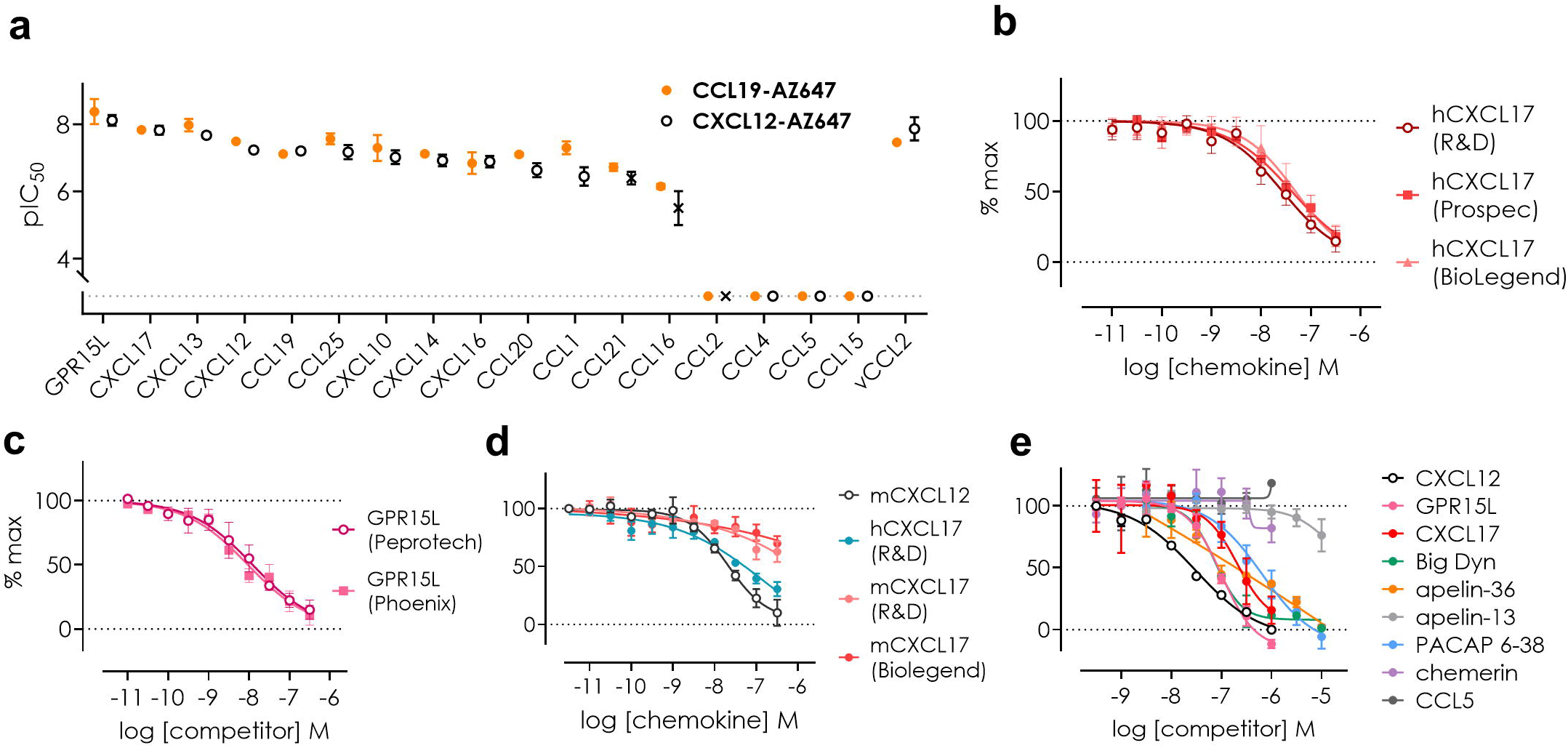
Confirmation of GPR182 ligands binding. **(a)** pIC50 values for a selected panel of human chemokines determined by flow cytometry binding competition with either AZ647-labelled CXCL12 (10 nM) or AZ647-labelled CCL19 (20 nM). **(b-d)** Concentration–response curves of hCXCL17 (b), hGPR15L (c) and mCXCL17 (d) from distinct suppliers determined by flow cytometry binding competition with human or mouse AZ647-labelled CXCL12 (10 nM) on GPR182. **(e)** Concentration– response curves of selected ligands from different families determined by NanoBRET binding competition with AZ488-labelled CXCL12 (10 nM) on purified membranes from HEK293-GPR182. Results are expressed as mean ± SEM of at least three independent experiments.

**Fig. S3.**
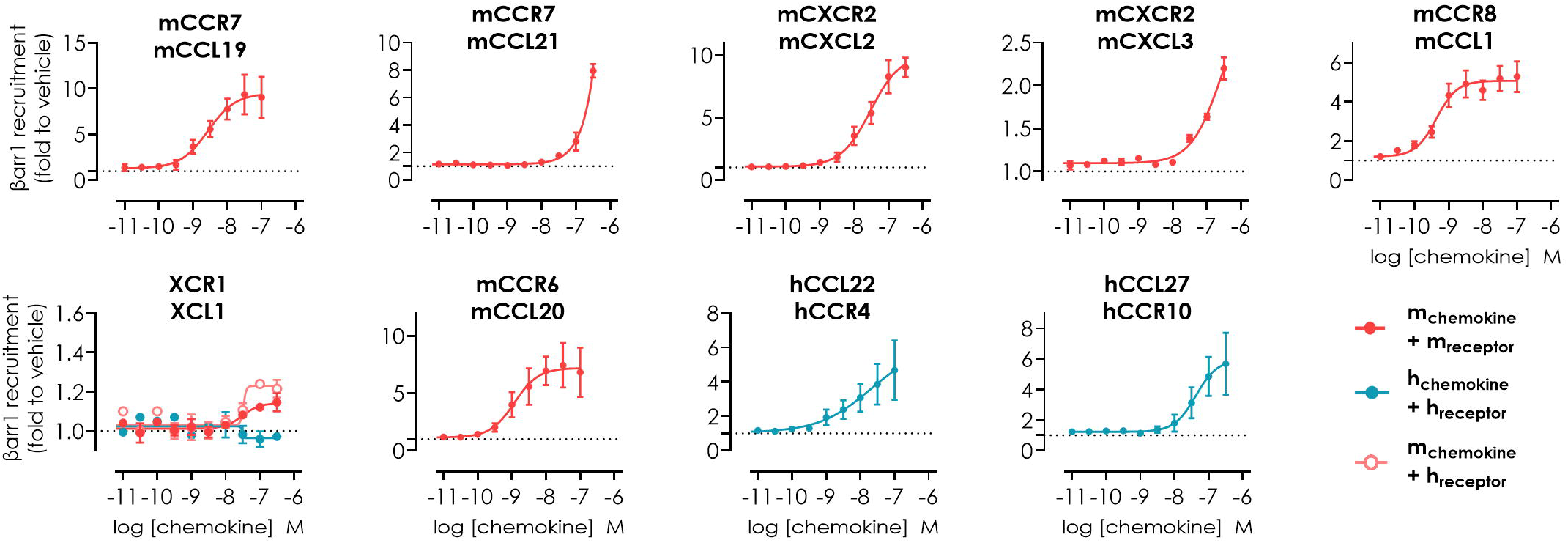
Verification of the activity of chemokines of interest. β-arrestin 1 recruitment to the cognate receptor of chemokines (human or mouse) for which only one of the two species orthologues was active on GPR182 as presented in Figure 1a. This confirms the functionality of these chemokines towards their cognate receptors. Results are expressed as mean ± SEM of at least three independent experiments.

**Fig. S4.**
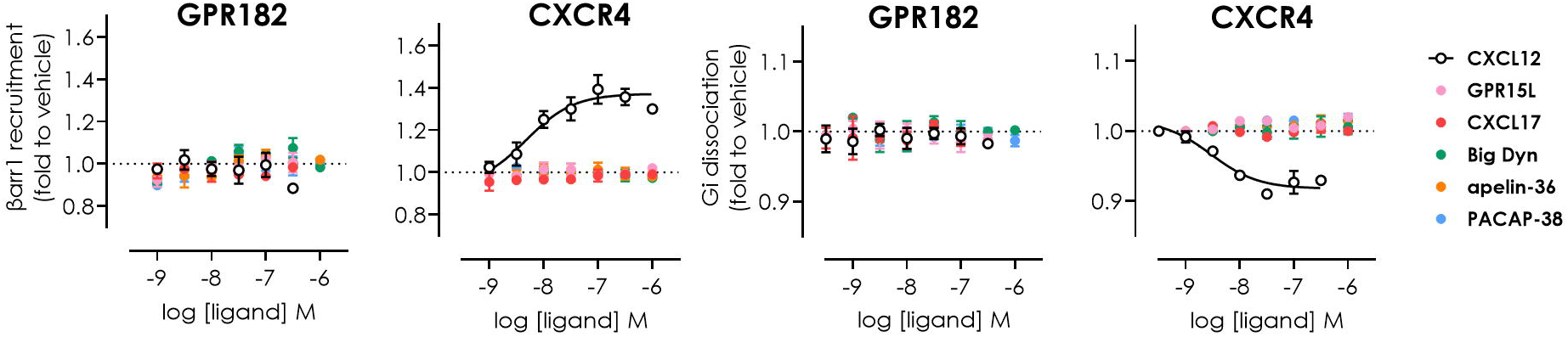
β-arrestin interaction and G protein activation following ligand binding to GPR182. NanoBiT β-arrestin 1 recruitment (left) and NanoBRET Gα_i_ protein dissociation (right) following binding of selected ligands to GPR182. CXCR4 was used as positive control. Results are expressed as fold to vehicle and mean ± SEM of at least three independent experiments.

**Fig. S5.**
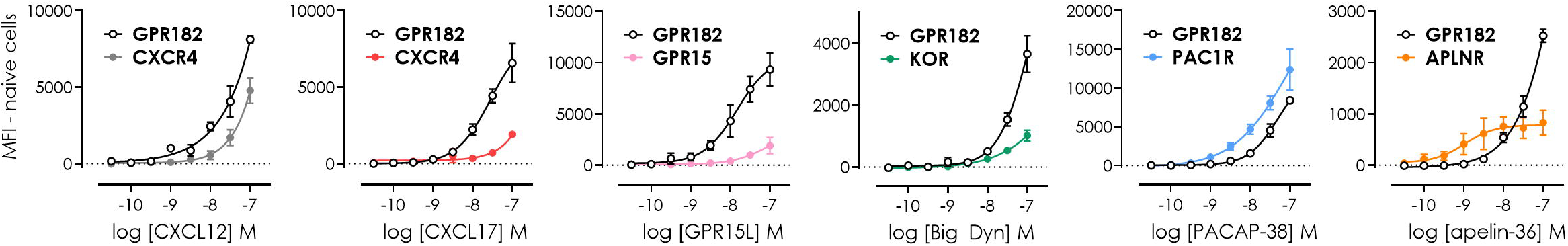
Uptake of Cy5-labelled ligands by GPR182- or cognate receptor expressing cells. Concentration-response curves of the uptake of Cy5-labelled chemokines or peptides by cells expressing mNeonGreen-tagged receptors measured by flow cytometry. For each ligand, the MFI from mNeonGreen-expressing cells was normalized on the signal obtained from naive HEK293 cells incubated with the respective ligand. Results are expressed as mean ± SEM of at least three independent experiments.

**Fig. S6.**
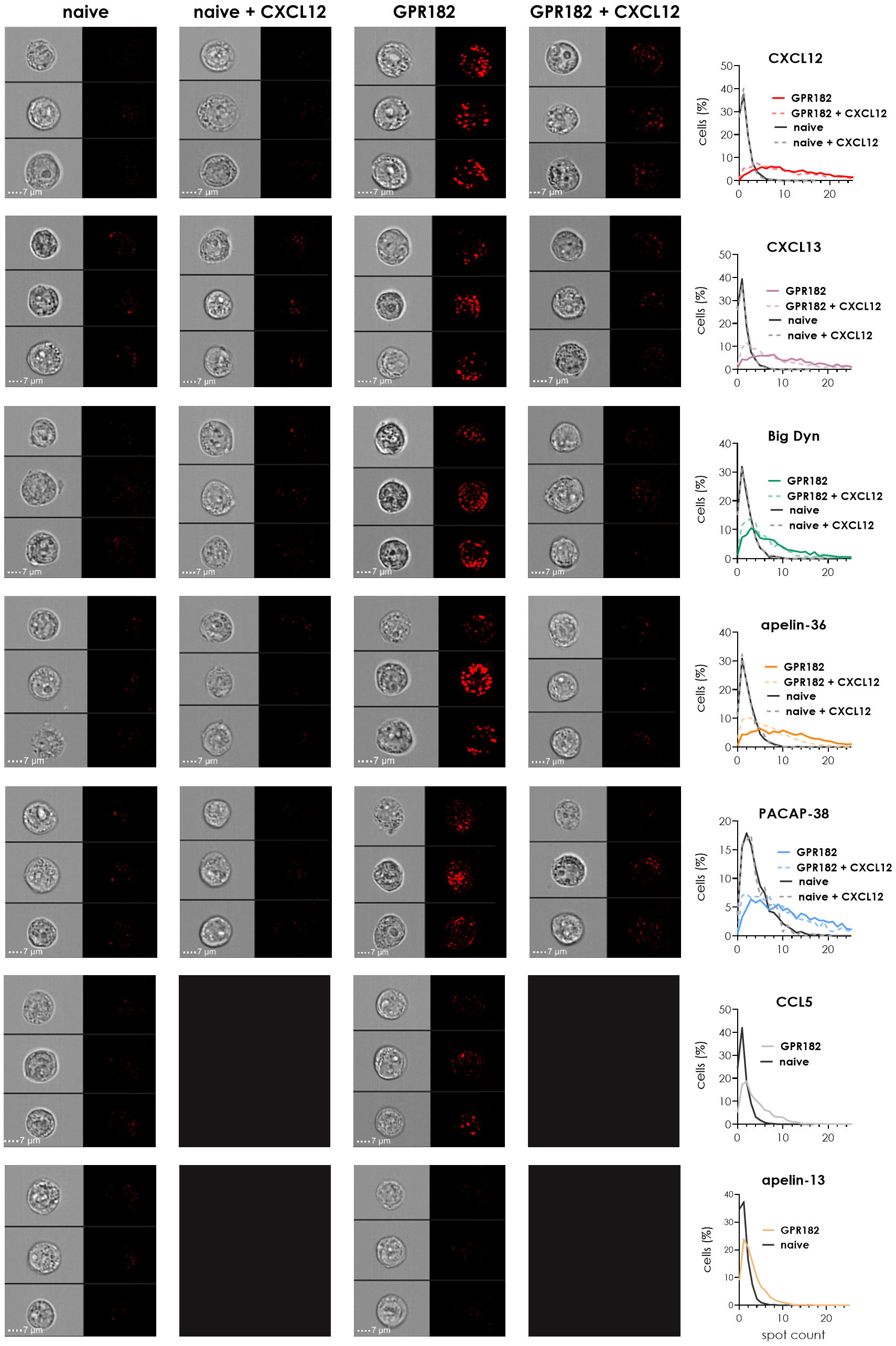
Uptake of Cy5-labelled ligands by GPR182 expressing cells in imaging flow cytometry. Uptake of Cy5-labelled CXCL12, CXCL13, PACAP-38, big dynorphin, apelin-36, CCL5 and apelin-13 (chemokines 10 nM, peptides 20 nM) in presence or absence of competing unlabelled CXCL12 (300 nM) by either naive HEK293 cells or cells expressing GPR182. (*left*) Representative images from three independent experiments acquired by imaging flow cytometry. Scale bar: 7 µM. (*right*) Corresponding percentage of cells with a given number of distinguishable vesicle-like structures (spots).

**Fig. S7.**
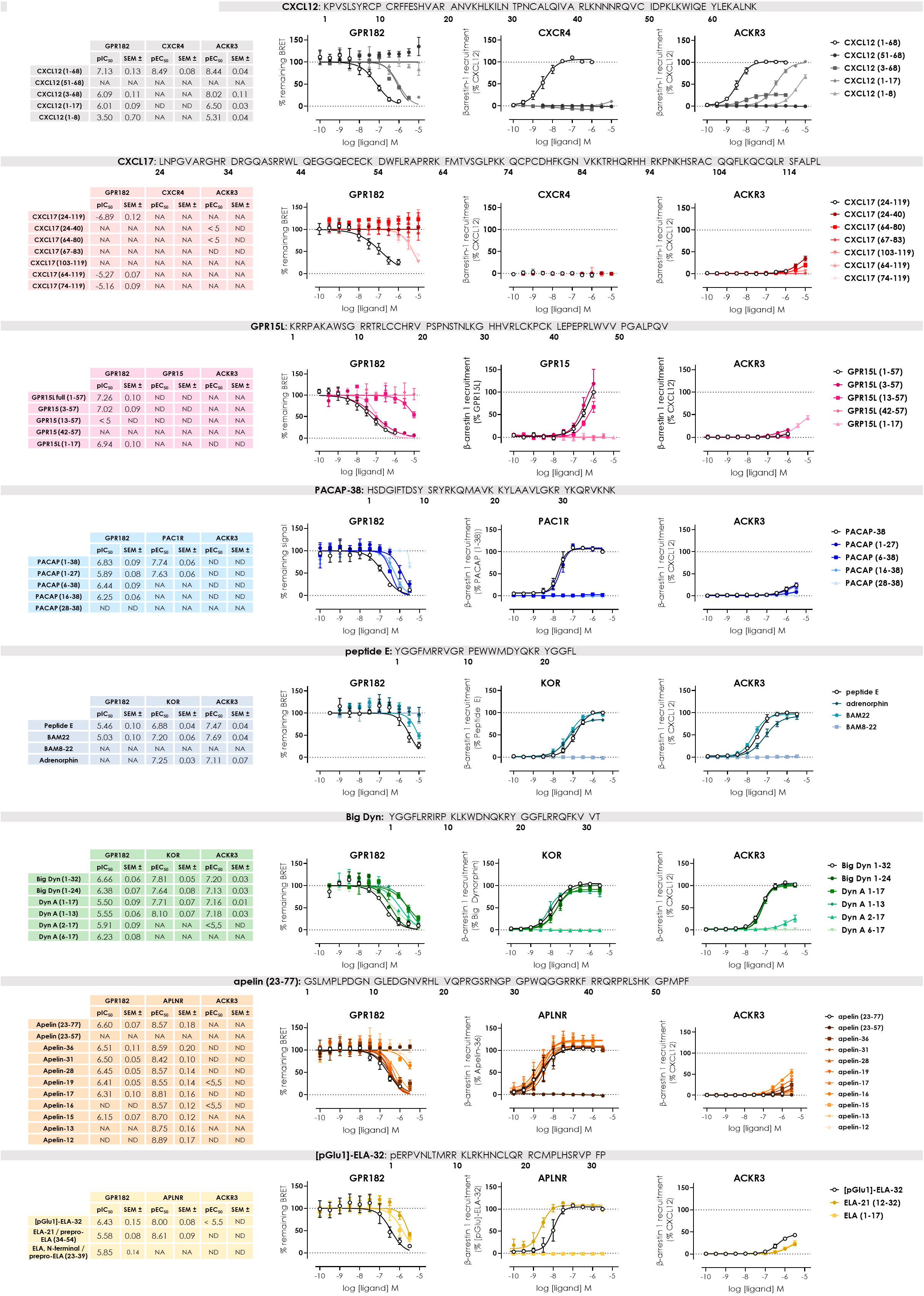
Effect of N- and C-terminal truncations in discovered ligands on their binding to GPR182 or the activation of their cognate receptor and ACKR3. Concentration–response curves of each ligand family determined by NanoBRET binding competition with AZ568-labelled CXCL12 (30 nM) on GPR182 (*left graph*) or ligand-induced β-arrestin 1 recruitment towards the cognate receptor (*middle*) or ACKR3 (*right*). pIC_50_ or pEC_50_ values are depicted in a table left to each peptide family. Sequences for the respective full-length ligands are indicated on top of each panel.

**Fig. S8.**
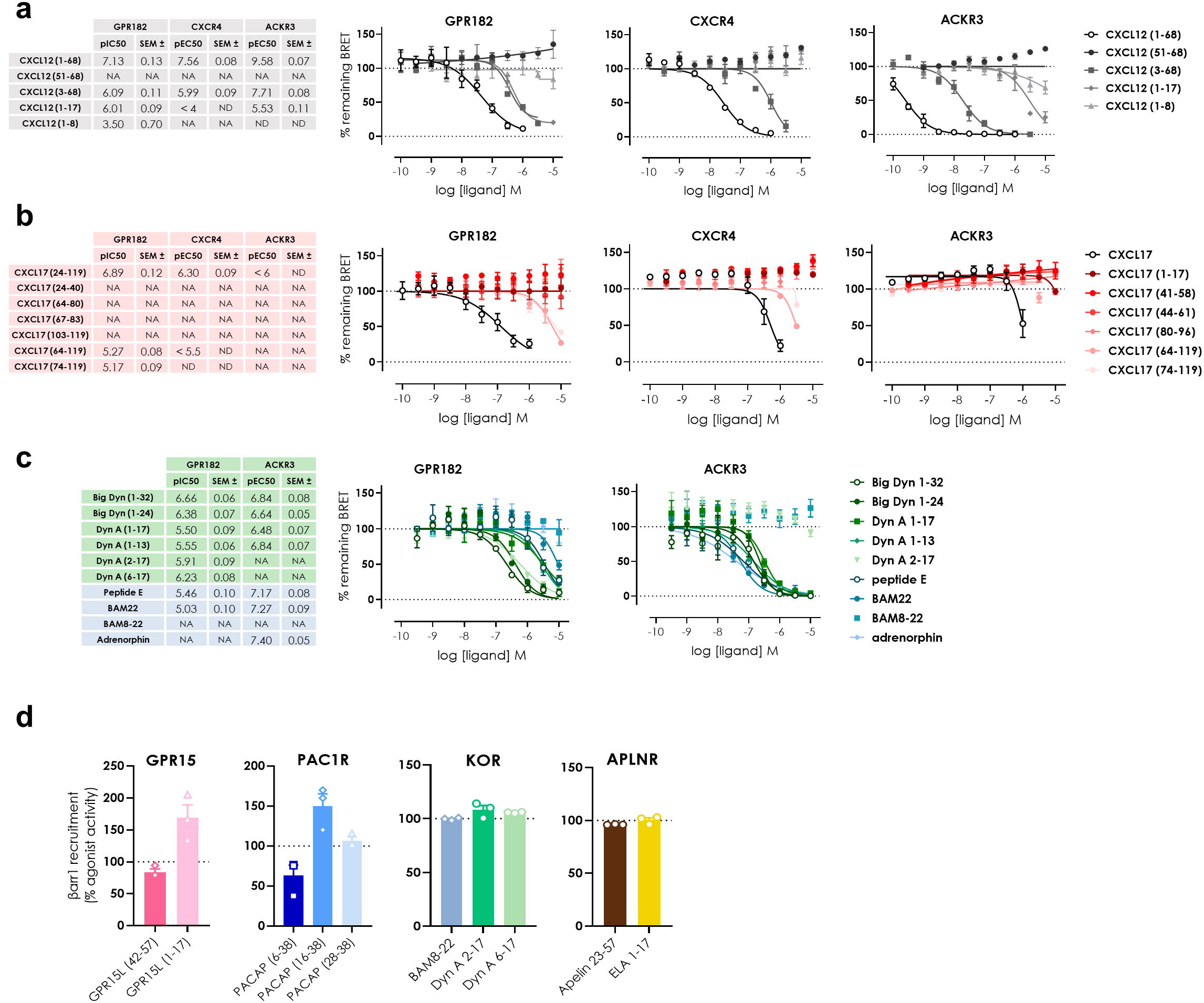
Assessment of binding and inhibitory activity of truncated fragments derived from the newly identified GPR182 ligands. **(a, b and c)** Concentration–response curves of full length and truncated forms of CXCL12 (a), CXCL17 (b) or opioid peptides (c) determined by NanoBRET binding competition. Competition was performed against AZ568-labelled CXCL12 (30 nM for GPR182 and CXCR4 or 2 nM CXCL12 for ACKR3). **(d)** Antagonist activity of the truncated peptides (3 µM) was measured through a β-arrestin 1 recruitment NanoBiT assay, following addition of GPR15L (300 nM) on GPR15, PACAP-38 (30 nM) on PAC1R, big dynorphin (30 nM) on KOR or apelin-15 (10 nM) on APLNR, in order to cover all potential binders. Results are expressed as percentage of signal received from stimulation with cognate ligands alone. For all panels, results are expressed as mean ± SEM of at least three independent experiments.

**Fig. S9.**
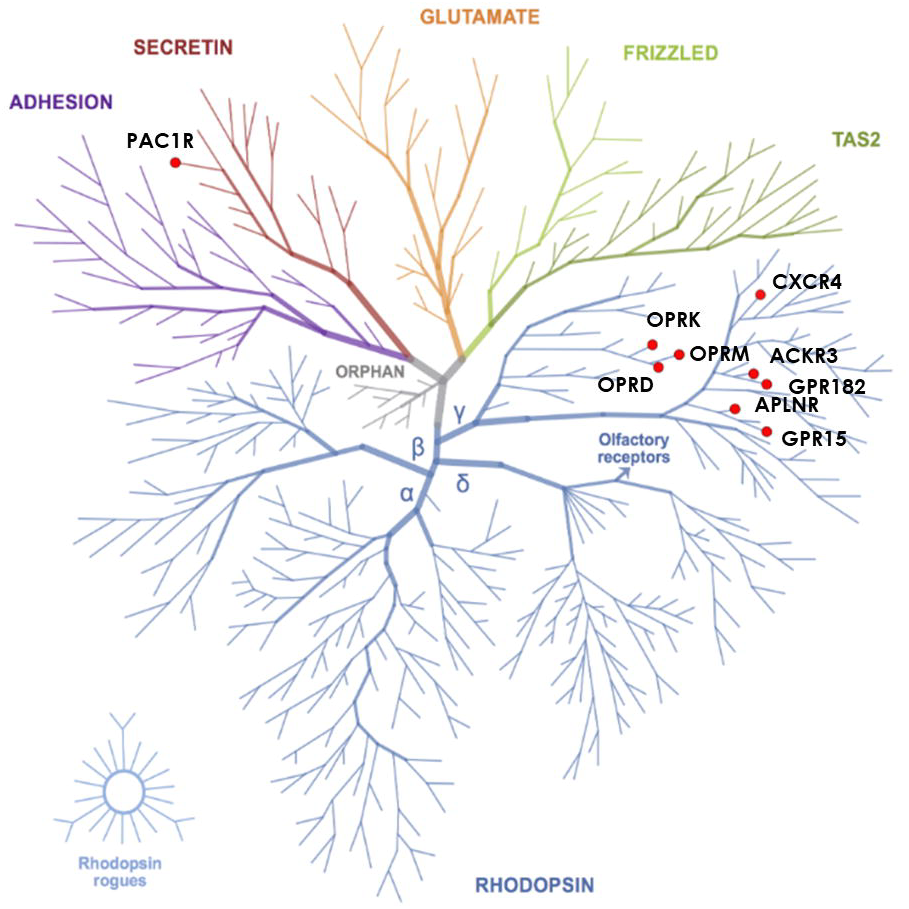
GPCR phylogenetic tree localizing GPR182, ACKR3 and the known receptors for the newly identified ligands of GPR182. This tree was created using the GPCR Tree Mapper (https://klifs.net/gpcr_mapper/)

